# The central amygdala controls learning in the lateral amygdala

**DOI:** 10.1101/126649

**Authors:** Kai Yu, Sandra Ahrens, Xian Zhang, Hillary Schiff, Charu Ramakrishnan, Lief Fenno, Karl Deisseroth, Pengcheng Zhou, Liam Paninski, Bo Li

**Author notes:** These authors contributed equally to this work (Sandra Ahrens, Xian Zhang). Correspondence to: Bo Li **Lead Contact:** Bo Li.

## Abstract

Both the lateral and the central nuclei of the amygdala are required for adaptive behavioral responses to environmental cues predicting threats. While experience-driven synaptic plasticity in the lateral amygdala is thought to underlie the formation of association between a sensory stimulus and an ensuing threat, how the central amygdala participates in such learning process remains unclear. Here we show that a specific class of central amygdala neurons, the protein kinase C-δ-expressing neurons, is essential for the synaptic plasticity underlying learning in the lateral amygdala, as it is required for lateral amygdala neurons to respond to unconditioned stimulus, and furthermore carries information about the unconditioned stimulus to instruct learning. Our results uncover an amygdala functional organization that may play a key role in diverse learning processes.

## INTRODUCTION

Adaptive behavioral responses to a threat are dependent on memories linking the threat with its associated environmental cues. Extensive evidence indicates that such memories are formed in the lateral amygdala (LA), in which coincident occurrences of a neutral environmental cue (also known as the conditioned stimulus (CS)) and a threatening event (also known as the unconditioned stimulus (US)), as exemplified in Pavlovian auditory fear conditioning (FC) by pairings of a sound with electrical footshock, induce Hebbian plasticity (Duvarci and Pare, 2014; Grundemann and Luthi, 2015; Herry and Johansen, 2014; LeDoux, 2000). This plasticity, expressed as strengthening of the synapses onto LA neurons driven by inputs carrying CS information, is associated with enhanced LA neuron responsiveness to the CS, required for the acquisition and recall of conditioned defensive responses, and thus, considered as a cellular substrate of aversive memory (Duvarci and Pare, 2014; Grundemann and Luthi, 2015; Herry and Johansen, 2014; LeDoux, 2000).

Recent studies demonstrate that the central amygdala (CeA), specifically its lateral division (CeL), is another amygdala nucleus indispensable for learning during fear conditioning (Ciocchi et al., 2010; Goosens and Maren, 2003; Li et al., 2013; Penzo et al., 2014; Penzo et al., 2015; Wilensky et al., 2006; Yu et al., 2016). Other studies also implicate the CeA in cognitive functions, such as attention (Calu et al., 2010; Lee et al., 2011; Roesch et al., 2012), that are critical for learning. Nevertheless, how the CeL contributes to the learning process that links environmental cues with threats remains poorly understood.

In traditional views of the fear circuit, the LA and the CeA are the main input and output, respectively, nuclei of the amygdala, so that information flows from the LA to the CeA (Herry and Johansen, 2014; LeDoux, 2007; Wilensky et al., 2006). Surprisingly, direct evidence supporting such serial information processing in FC has been lacking. On the other hand, previous studies have described functions of the CeA – including its involvement in the attention or alerting processes – that are independent of the LA during reinforcement learning (Balleine and Killcross, 2006; Haney et al., 2010; Roesch et al., 2012), suggesting that the two nuclei are not simply organized in series. In this study, we used a combination of electrophysiological, molecular, optogenetic, chemogenetic, and *in vivo* calcium imaging techniques to examine the functional relationship between distinct classes of CeA neurons and LA neurons in FC.

## RESULTS

### Somatostatin-expressing CeL neurons are not required for the FC-induced LA synaptic plasticity

We reasoned that, if information indeed flows serially from the LA to the CeA during FC, then inhibiting the CeA should leave the FC-induced LA synaptic plasticity intact. To test this hypothesis, we set out to inhibit the major classes of CeA neurons in mice with the tetanus toxin light chain (TeLC), which blocks neurotransmitter release (Murray et al., 2011). We first targeted somatostatin-expressing (^SOM+^) CeL neurons by bilaterally injecting the CeL of *Som-Cre* mice, in which the Cre recombinase is expressed under the endogenous Som promoter (Taniguchi et al., 2011), with an adeno-associated virus expressing the TeLC-GFP or GFP (as a control) in a Cre-dependent manner (AAV-DIO-TeLC-GFP or AAV-DIO-GFP, respectively) (Murray et al., 2011) (Fig. 1A-F). In the same mice we also injected the auditory thalamus (the medial geniculate nucleus, MGN), which transmits CS information in auditory FC, with an AAV expressing the light-gated cation channel channelrhodopsin (AAV-CAG-ChR2(H134R)-eYFP) (Zhang et al., 2006) (Fig. 1A, F).

**Fig. 1.**
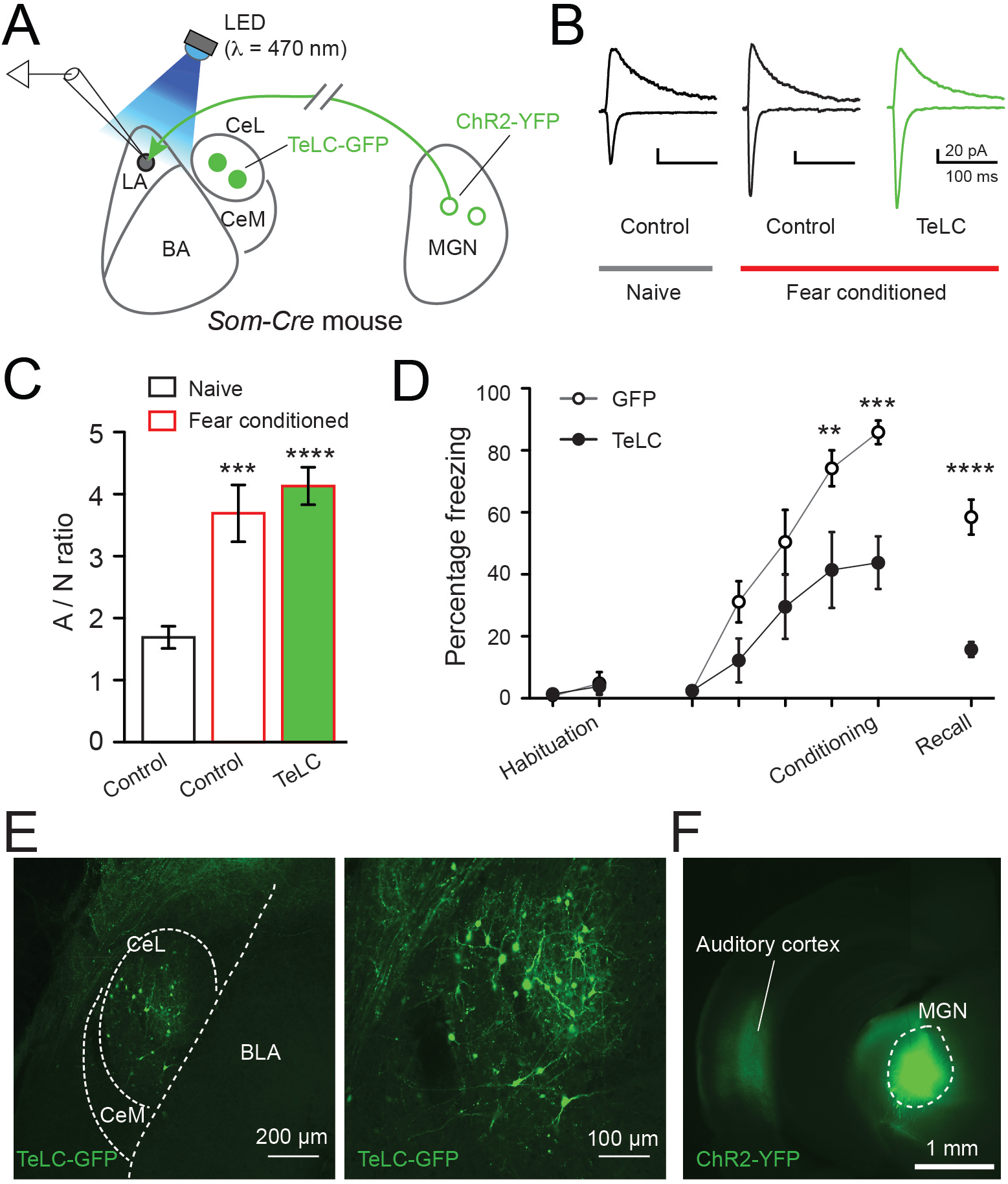
SOM^+^ CeL neurons are not required for plasticity underlying learning in the LA. (A) A schematic of the experimental configuration. (B) Example traces of the excitatory postsynaptic currents (EPSCs). The outward currents were recorded at a holding potential of +40 mV, and the inward currents were recorded at −70 mV. (C) Quantification of the ratio between the average amplitude of EPSCs mediated by AMPA receptors and that mediated by NMDA receptors (A/N) (from left to right: n = 19 cells/3 mice, 15 cells/3 mice, and 26 cells/4 mice; F(2,57) = 17.2; ***P < 0.001, ****P < 0.0001 compared with the control naïve group; one-way ANOVA followed by Bonferroni’s test). (D) Quantification of freezing behaviour (GFP, n = 5, TeLC, n = 5, F(1,8) = 21.06, ****P < 0.0001, ***P < 0.001, **P < 0.01, two-way RM-ANOVA followed by Bonferroni’s test). (E) Images of a coronal brain section from a representative mouse, in which the SOM^+^ CeL neurons expressed TeLC-GFP. The image on the right is a higher magnification of the CeL area in the image on the left. (F) Image of a coronal brain section from the same mouse as that in (E), showing the expression of ChR2-YFP in the MGN, which gave rise to axons in the auditory cortex. Data are presented as mean ± s.e.m. in C, D.

After viral expression had reached optimal levels, we subjected the mice to auditory FC followed by preparation of acute brain slices, in which we recorded from LA neurons the AMPA receptor (A) and the NMDA receptor (N) mediated components of synaptic responses evoked by light-stimulation of MGN axons (Fig. 1A-C). As expected, inhibition of SOM^+^ CeL neurons did not affect the FC-induced synaptic strengthening, measured as an increase in A/N ratio (Clem and Huganir, 2010; Nabavi et al., 2014), onto LA neurons (Fig. 1B, C). This manipulation did cause impairment in conditioned freezing behavior – a characteristic defensive response in rodents (LeDoux et al., 1988) (Fig. 1D) – an effect that is consistent with our previous findings (Li et al., 2013; Penzo et al., 2015).

### Protein kinase C-δ-expressing CeL neurons are required for the FC-induced LA synaptic plasticity and learning

We next examined the effects of inhibiting protein kinase C-δ-expressing (PKC-δ^+^) CeL neurons, another major population in the CeL, with the TeLC in *Prkcd-Cre* mice that express Cre in PKC-δ^+^ CeL neurons (Haubensak et al., 2010) (Fig. 2A-D, Fig. S1A-C). To our surprise, inhibiting PKC-δ^+^ CeL neurons completely abolished the FC-induced synaptic strengthening onto LA neurons (Fig. 2A-C). Notably, unilateral inhibition of PKC-δ^+^ CeL neurons was sufficient to abolish the synaptic strengthening in the ipsilateral LA, while leaving that in the contralateral LA intact (Fig. 2B, C). Furthermore, bilateral inhibition of PKC-δ^+^ CeL neurons drastically impaired conditioned freezing (Fig. 2D; Fig. S1A, B), while unilateral inhibition of these neurons was less effective (Fig. S1A). Bilateral inhibition of PKC-δ^+^ CeL neurons also completely abolished conditioned lick-suppression, wherein a sound predicting tail shock suppresses licking for water rewards (Yu et al., 2016) (Fig. 3) (see Methods). Such conditioned suppression of action, like the conditioned freezing, has been shown to depend on both LA plasticity (Erlich et al., 2012; Nabavi et al., 2014) and CeA function (Killcross et al., 1997; Yu et al., 2016).

**Fig. 2.**
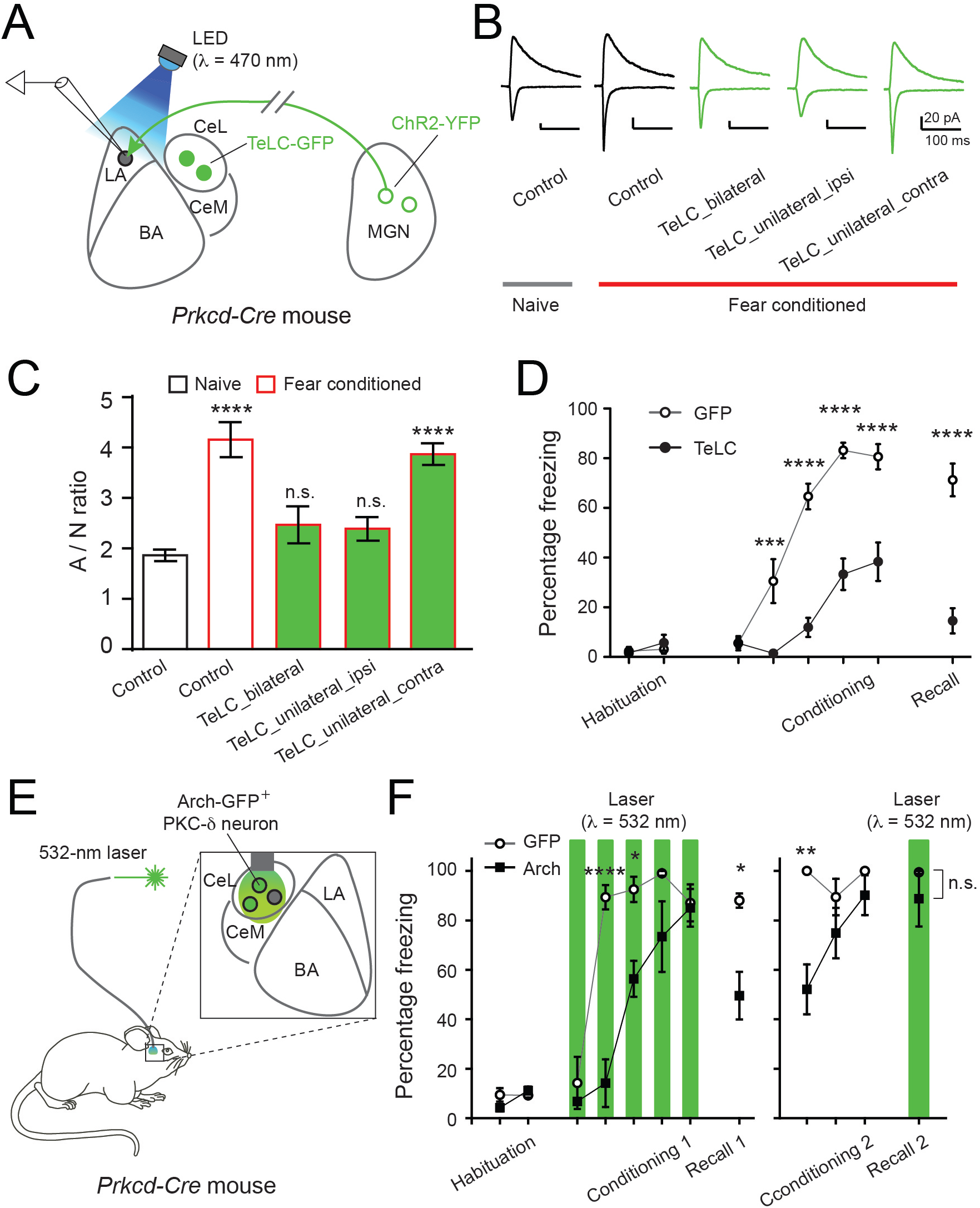
PKC-δ^+^ CeL neurons are required for plasticity underlying learning in the LA. (A) A schematic of the experimental configuration. (B) Example traces of the excitatory postsynaptic currents (EPSCs). (C) Quantification of A/N (from left to right: n = 20 cells/3 mice, 22 cells/3 mice, 9 cells/2 mice, 29 cells/4 mice, 30 cells/4 mice; F(4,105) = 15.28, P < 0.0001; ****P < 0.0001, n.s., not significant (P > 0.05), compared with the control naïve group; one-way ANOVA followed by Bonferroni’s test). (D) Quantification of freezing behaviour (GFP, n = 11, TeLC, n = 11, F(1,20) = 57.88, P < 0.0001; ****P < 0.0001, ***P < 0.001, two-way RM-ANOVA followed by Bonferroni’s test). (E) A schematic of the experimental configuration. (F) Quantification of freezing behaviour (GFP, n = 3, Arch, n = 4, F(1,5) = 52.41, P < 0.001; *P < 0.05, **P < 0.01, ****P < 0.0001, n.s., not significant (P > 0.05); two-way RM-ANOVA followed by Bonferroni’s test). Data are presented as mean ± s.e.m. in C, D, and F.

**Fig. 3.**
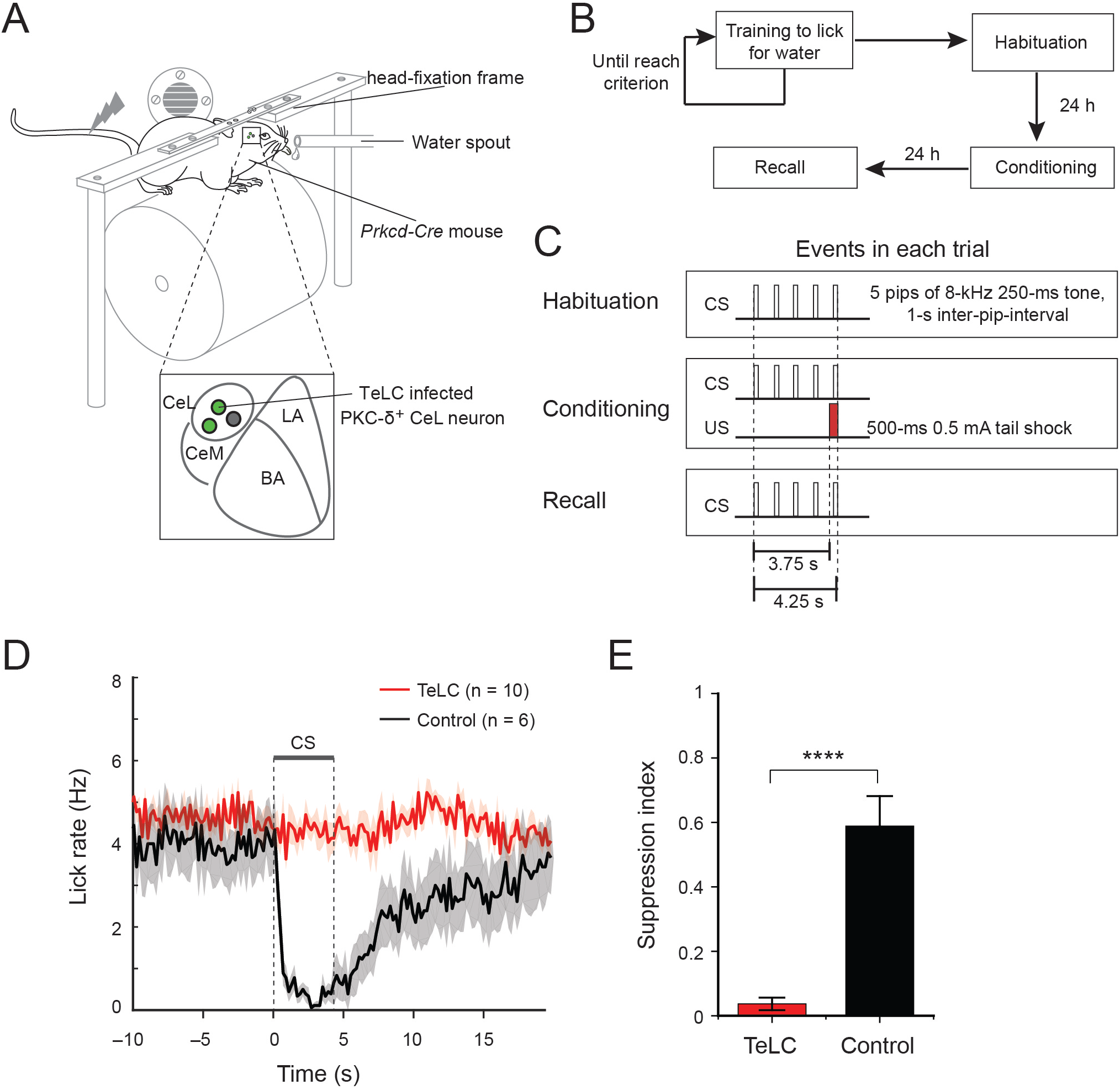
PKC-δ^+^ CeL neurons are required for conditioned lick suppression. (A) A schematic of the experimental configuration. (B & C) Schematics of the experimental procedure for training mice in the conditioned lick suppression task. (D) Lick rate for mice during recall. Shaded areas represent s.e.m. (E) Conditioned lick suppression, quantified as suppression index (TeLC, n = 10, control, n = 6, T(14) = 7.36, ****P < 0.0001, t test). Data are presented as mean ± s.e.m. in D, E.

The behavioral effect of PKC-δ^+^ CeL neuron inhibition is likely caused by impairment in learning rather than expression of the defensive responses, as suggested by the impaired LA synaptic plasticity. To verify this possibility we used optogenetics, for which we bilaterally injected the CeL of the *Prkcd-Cre* mice with a Cre-dependent AAV expressing the light-sensitive proton pump archaerhodopsin (Chow et al., 2010) (AAV-DIO-Arch-GFP), and subsequently implanted optical fibres above the CeL for light delivery (Fig. 2E, Fig. S1D, E). Optogenetic inhibition of PKC-δ^+^ CeL neurons during conditioning, but not during memory recall, significantly reduced conditioned freezing behavior (Fig. 2F). Together, these results indicate that the activity of PKC-δ^+^ CeL neurons is required for both the FC-induced LA synaptic plasticity and learning.

Of note, it has been proposed that PKC-δ^+^ CeL neurons have a different function, such that inhibition of these neurons may cause freezing through disinhibition of the medial division of the CeA (CeM) (Haubensak et al., 2010). However, we found that optogenetic inhibition of PKC-δ^+^ CeL neurons failed to induce any freezing behavior or aversive response in a real time place aversion (RTPA) task in naïve mice (Fig. S2).

### PKC-δ^+^ CeL neurons develop CS responses after fear conditioning

Why are neurons in the CeL required for synaptic plasticity and learning in the LA? In auditory FC, the coincident detection of sound (CS) and shock (US) by LA neurons is thought to be a prerequisite for these neurons to undergo the synaptic strengthening underlying learning. While sound can reach the LA via the MGN and the auditory cortex, the route through which shock is transmitted to the LA remains unclear (Herry and Johansen, 2014). Interestingly, it has recently been shown that CeL neurons, including PKC-δ^+^ neurons, are the direct postsynaptic targets of the parabrachial nucleus (PBN) (Han et al., 2015), a brainstem structure that provides nociceptive signals, and that activation of the PBN–CeL pathway is sufficient to drive aversive learning (Han et al., 2015; Sato et al., 2015). These findings raise the possibility that PKC-δ^+^ CeL neurons may participate in conveying US information during FC.

To test this possibility, we examined whether and how PKC-δ^+^ CeL neurons might respond to US. We delivered GCaMP6, the genetically encoded calcium indicator (Chen et al., 2013), into PKC-δ^+^ neurons by injecting the CeL of the *Prkcd-Cre;Som-Flp* mice with an intersectional AAV-C_on_/F_off_-GCaMP6m (Fenno et al., 2014). This strategy ensures specific infection of PKC-δ^+^ neurons and avoids infection of a small fraction of CeL neurons expressing both PKC-δ and SOM (Li et al., 2013). The same mice were implanted in the CeL with gradient-index (GRIN) lenses, through which the calcium signals could be recorded using a miniature integrated fluorescence microscope (Fig. 4A; see Methods) (Ghosh et al., 2011; Jennings et al., 2015; Resendez et al., 2016; Ziv et al., 2013). We subsequently trained these mice in the conditioned lick-suppression task (Fig. 3) (Yu et al., 2016) while imaging PKC-δ^+^ CeL neuron calcium responses (Fig. 4A; Fig. S3).

**Fig. 4.**
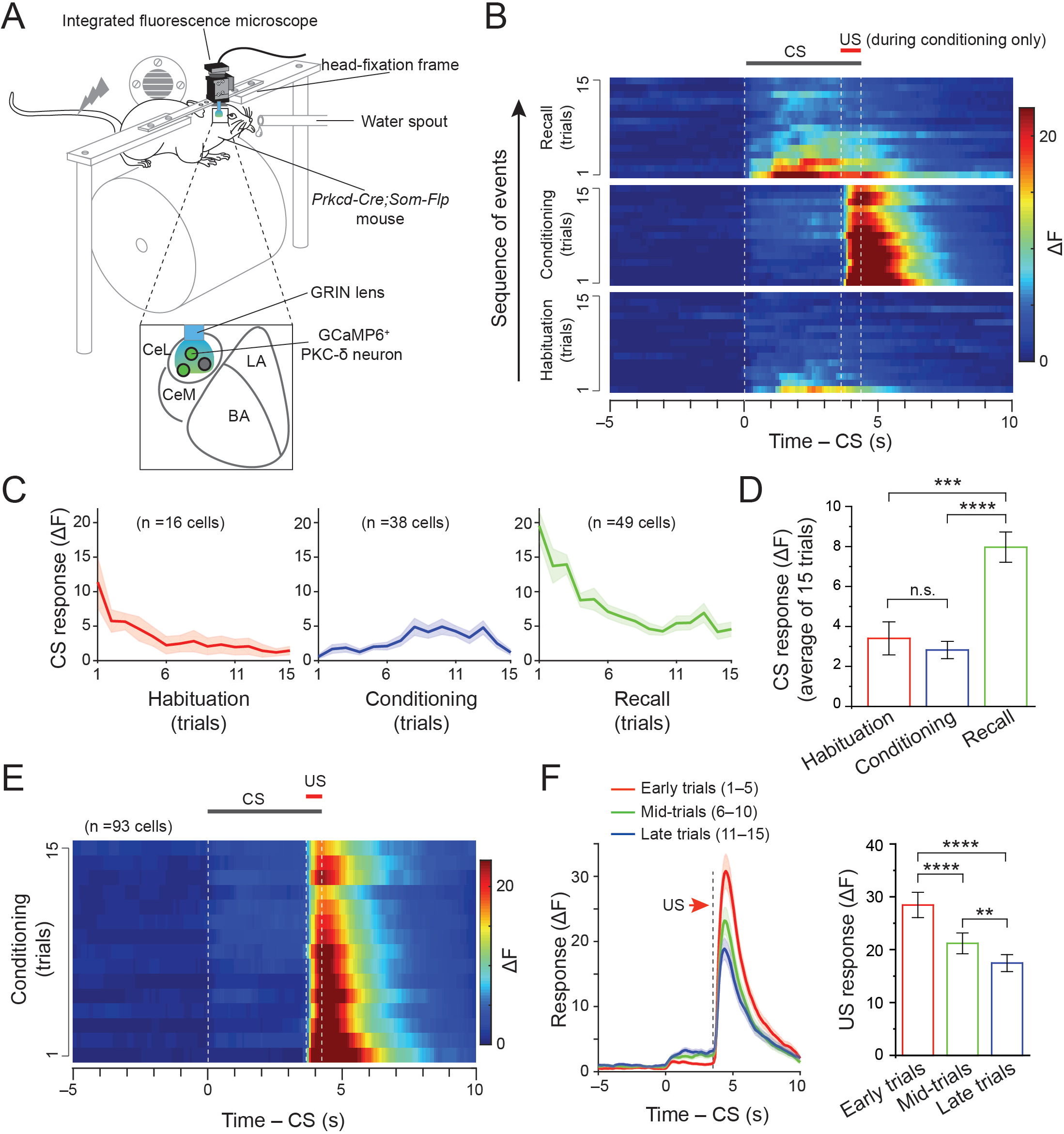
The CS and US responses in PKC-δ^+^ CeL neurons during fear conditioning. (A) A schematic of the experimental configuration. (B) Heat-maps of the average temporal calcium activities of all CS-responsive PKC-δ^+^ CeL neurons (data from 3 mice) for each trial during habituation, conditioning, and recall. Dashed lines indicate the timing of CS or US. (C) Average CS-induced responses in the same neurons as those in (B) for each trial. Shaded areas represent s.e.m. (D) CS responses of the same neurons as those in (C), averaged for all trials during habituation, conditioning, or recall (F(2,100) = 17.97, P < 0.0001; ***P < 0.001, ****P < 0.0001, n.s., not significant (P > 0.05); one-way ANOVA followed by Bonferroni’s test). (E) A heat-map of the average temporal calcium activities of all US-responsive PKC-δ^+^ CeL neurons for each trial during conditioning. Dashed lines indicate the timing of CS or US. (F) The time course (left) and peak amplitude (right) of US-evoked responses in the same neurons as those in (E), averaged for the trials in different stages of conditioning (n = 93 cells, 3 mice; F(1.5,140.2) = 26.41, P < 0.0001; **P < 0.01, ****P < 0.0001; one-way RM-ANOVA followed by Bonferroni’s test; shaded areas in (F) represent s.e.m.). Data are presented as mean ± s.e.m. in C, D, and F.

We identified active PKC-δ^+^ CeL neurons during habituation, conditioning and recall, based on the spontaneous activities and CS or US responses in these neurons (Fig. S3, S4; see Movie S1; and see Methods). The fractions of neurons showing CS-evoked responses were 10.2% (16/157), 20.0% (38/190), and 26.5% (49/185) during habituation, conditioning, and recall, respectively (P < 0.05, χ^2^ test with Bonferroni’s correction, comparing habituation with other groups) (Fig. 4B-D). On average, the CS responses of these neurons desensitized during habituation, but recovered quickly following US presentations during conditioning, and were enhanced during recall (Fig. 4B-D). These results indicate that PKC-δ^+^ CeL neurons gain CS responses following FC.

### PKC-δ^+^ CeL neurons show US responses that are modulated by expectation

During conditioning, 48.9% (93/190) of PKC-δ^+^ CeL neurons showed prominent US-evoked responses (Fig. 4E, F; Fig. S4). Interestingly, these responses decreased as conditioning progressed (Fig. 4E, F), consistent with theories and evidence that instructive US signals are suppressed when they become expected (Belova et al., 2007; Fanselow, 1998; Johansen et al., 2010; McNally et al., 2011; Ozawa et al., 2017; Rescorla and Wagner, 1972). To specifically test if expectation modulates the US responses, we tracked the responses of each PKC-δ^+^ CeL neuron to a series of US presentations, some of which were signaled by the CS while the others were delivered unexpectedly (Fig. 5A, B). We found that, among all the US-responsive neurons, 21.1% had stronger responses, while 8.8% showed weaker responses to unsignaled than to signaled US (Fig. 5B-E). These results demonstrate that about half of PKC-δ^+^ CeL neurons show robust US responses, with a subpopulation of these neurons having US responses suppressed by expectation, the property of a teaching signal according to theories of fear conditioning (McNally et al., 2011).

**Fig. 5.**
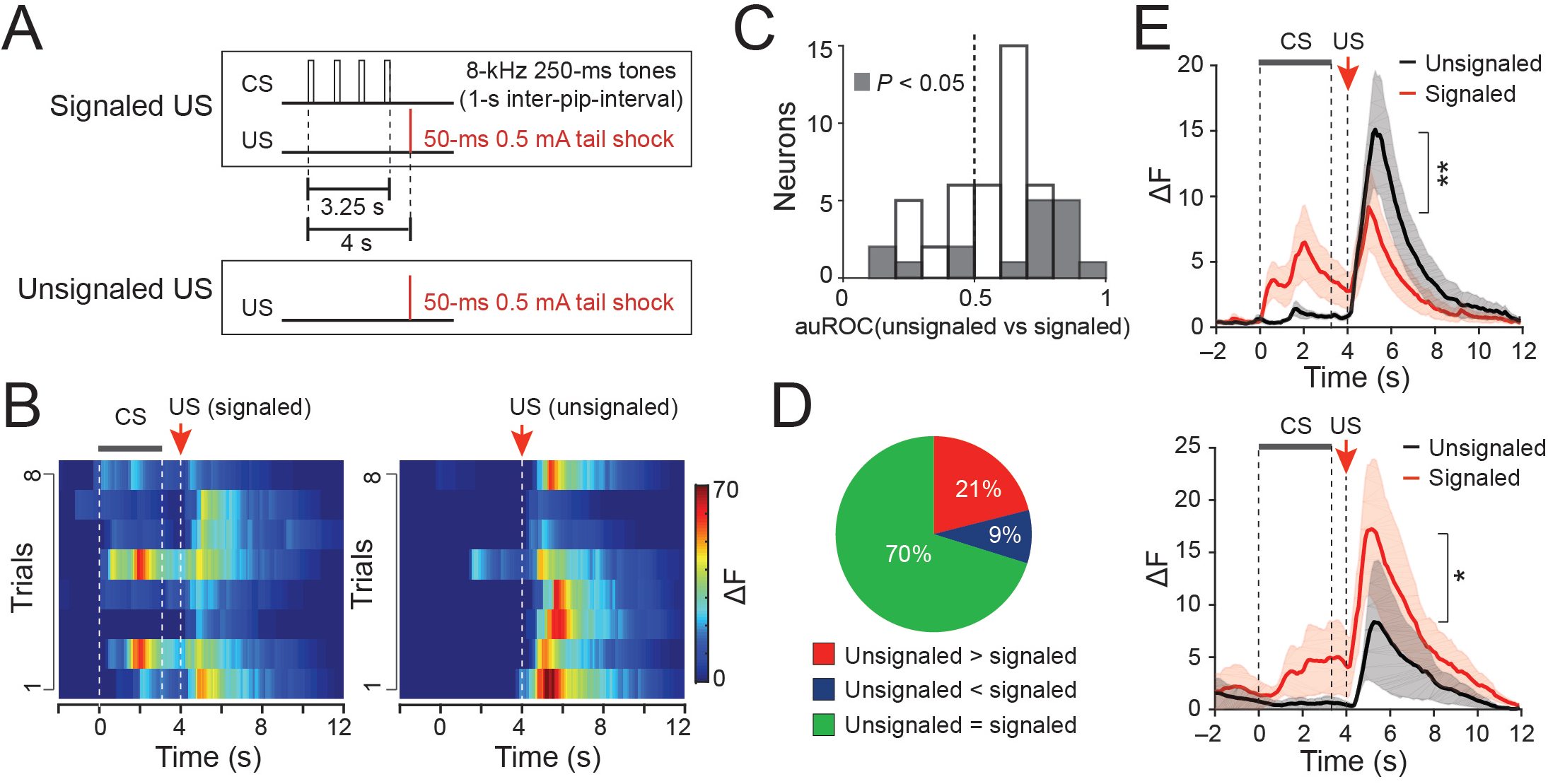
The US responses of PKC-δ^+^ CeL neurons are modulated by expectation. (A) Schematics for the delivery of signaled and unsignaled US. (B) Heat-maps of the temporal calcium activities of a representative PKC-δ^+^ CeL neuron, in trials where signaled (left) or unsignaled (right) US were delivered. Dashed lines indicate the timing of CS or US. (C) Results of auROC (area under the Receiver Operating Characteristic curve) analysis of differences between responses to unsignaled and signaled US. (D) Distribution of all US-responsive PKC-δ^+^ CeL neurons (57 cells, 4 mice) based on their response properties. (E) Average responses during unsignaled and signaled US trials, for PKC-δ^+^ CeL neurons that had stronger (upper panel) (n = 12, T(11) = 3.33, **P < 0.01, paired t test) or weaker (lower panel) (n = 5, T(4) = 3.96, *P < 0.05, paired t test) responses to unsignaled US than to signaled US. Data are presented as mean ± s.e.m. in E, with shaded areas representing s.e.m.

### PKC-δ^+^ CeL neuron activity is required for the US responses of LA neurons

We next examined the role of PKC-δ^+^ CeL neurons in conveying the US signal to the LA. We implanted GRIN lenses in the LA of *Prkcd-Cre* mice, in which the GCaMP6 was expressed in LA neurons by an AAV-hSyn-GCaMP6f, and in which an inhibitory DREADD (Designer Receptor Exclusively Activated by Designer Drug) molecule derived from the kappa-opioid receptor (KORD) (Fig. S5) (Vardy et al., 2015) was selectively expressed in PKC-δ^+^ CeL neurons by an AAV-hSyn-DIO-HA-KORD (Fig. 6A, Fig. S6). This strategy allowed us to track the US responses of the same LA neurons (see Fig. S7; Movie S2) before and after transiently inhibiting PKC-δ^+^ CeL neurons with systemically applied salvinorin B (SALB), the agonist of KORD (Vardy et al., 2015) (Fig. 6; Fig. S6, S7). Notably, we found that inhibition of PKC-δ^+^ CeL neurons significantly suppressed shock-evoked responses of LA neurons (Fig. 6B-E), indicating that PKC-δ^+^ CeL neurons play an important role in conveying the US signal to the LA.

**Fig. 6.**
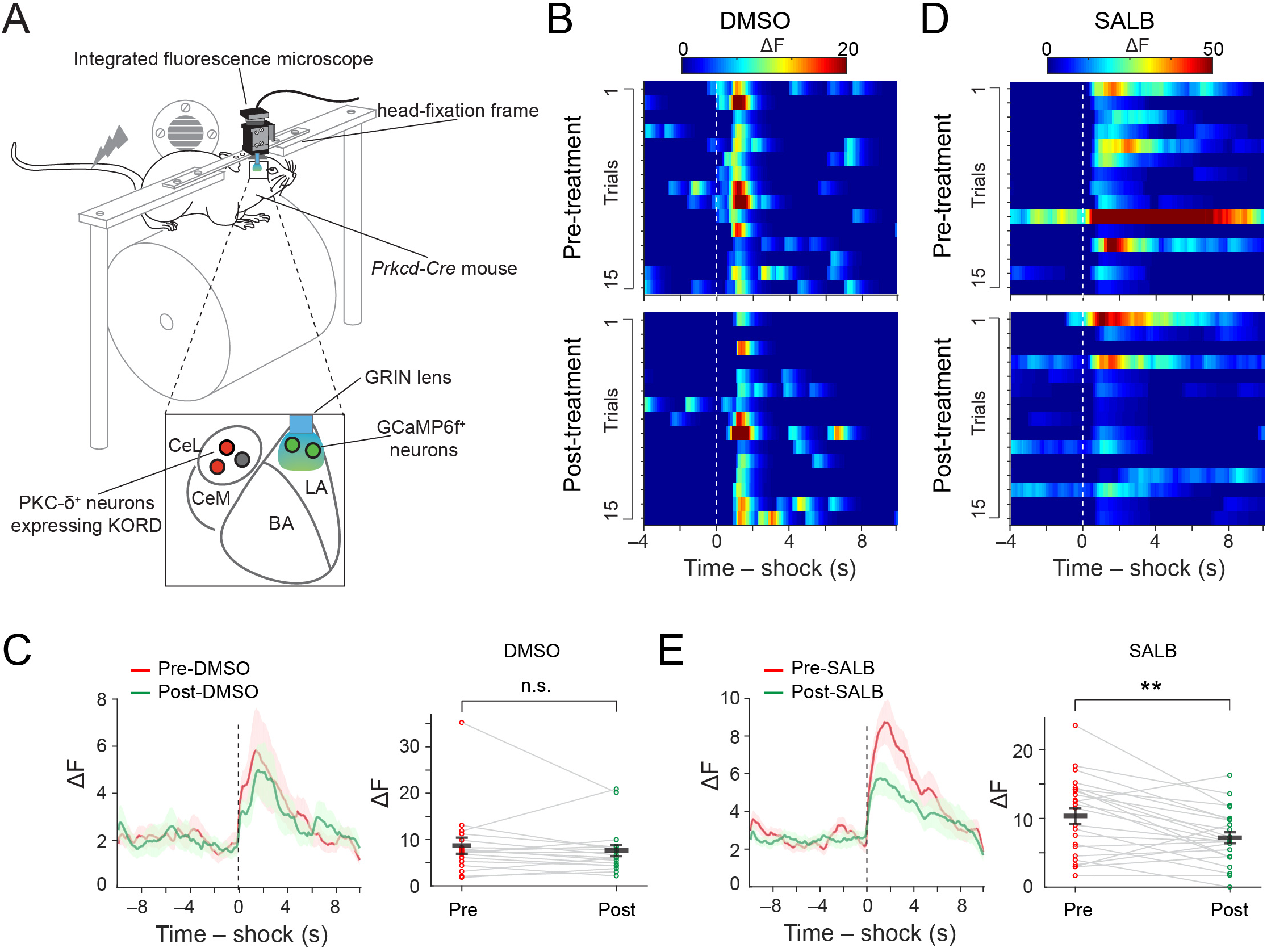
PKC-δ^+^ CeL neurons are required for the normal US responses of LA neurons. (A). A schematic of the experimental configuration. (B) Heat-maps of the temporal calcium activities of a representative LA neuron, before (top panel) and after (bottom panel) DMSO application. The dashed line indicates the onset of US. (C) Left: average temporal activities of all shock-responsive LA neurons aligned to shock onset (dashed line), before and after DMSO treatment. Shaded areas represent s.e.m. Right: scatter plot of the peak shock responses of each of the neurons before and after DMSO treatment (T(17) = 0.93, n.s., P > 0.05; paired t test; n = 18 neurons / 4 mice). (D & E) Same as in (B & C), except that SALB was applied instead of DMSO to the same mice in different imaging sessions (T(23) = 3.5, **P < 0.01; paired t test; n = 24 neurons / 4 mice). Data are presented as mean ± s.e.m. in C and E.

### PKC-δ^+^ CeL neurons convey aversive information and drive learning

If the US response of PKC-δ^+^ CeL neurons represents a teaching signal in FC, then it may also carry negative emotional valence and, moreover, be able to instruct learning. Indeed, we found that inhibition of these neurons with the TeLC reduced animals’ reaction to electrical shocks (Fig. 7A), suggesting that these neurons are important for processing the affective component of the US. Conversely, optogenetic activation of PKC-δ^+^ CeL neurons induced aversion in mice in the RTPA task (Fig. 7B, C; Fig. S8A), and caused marked lick suppression (Fig. 8A; Fig. S8A) (again, for the experiments in Fig. 7B, C and Fig. 8A, we injected the CeL of *Prkcd-Cre;Som-Flp* mice with an intersectional AAV-C_on_/F_off_-ChR2-mCherry (Fenno et al., 2014), to selectively infect PKC-δ^+^ neurons while avoid infecting the small fraction of CeL neurons expressing both PKC-δ and SOM (Li et al., 2013)).

**Fig. 7.**
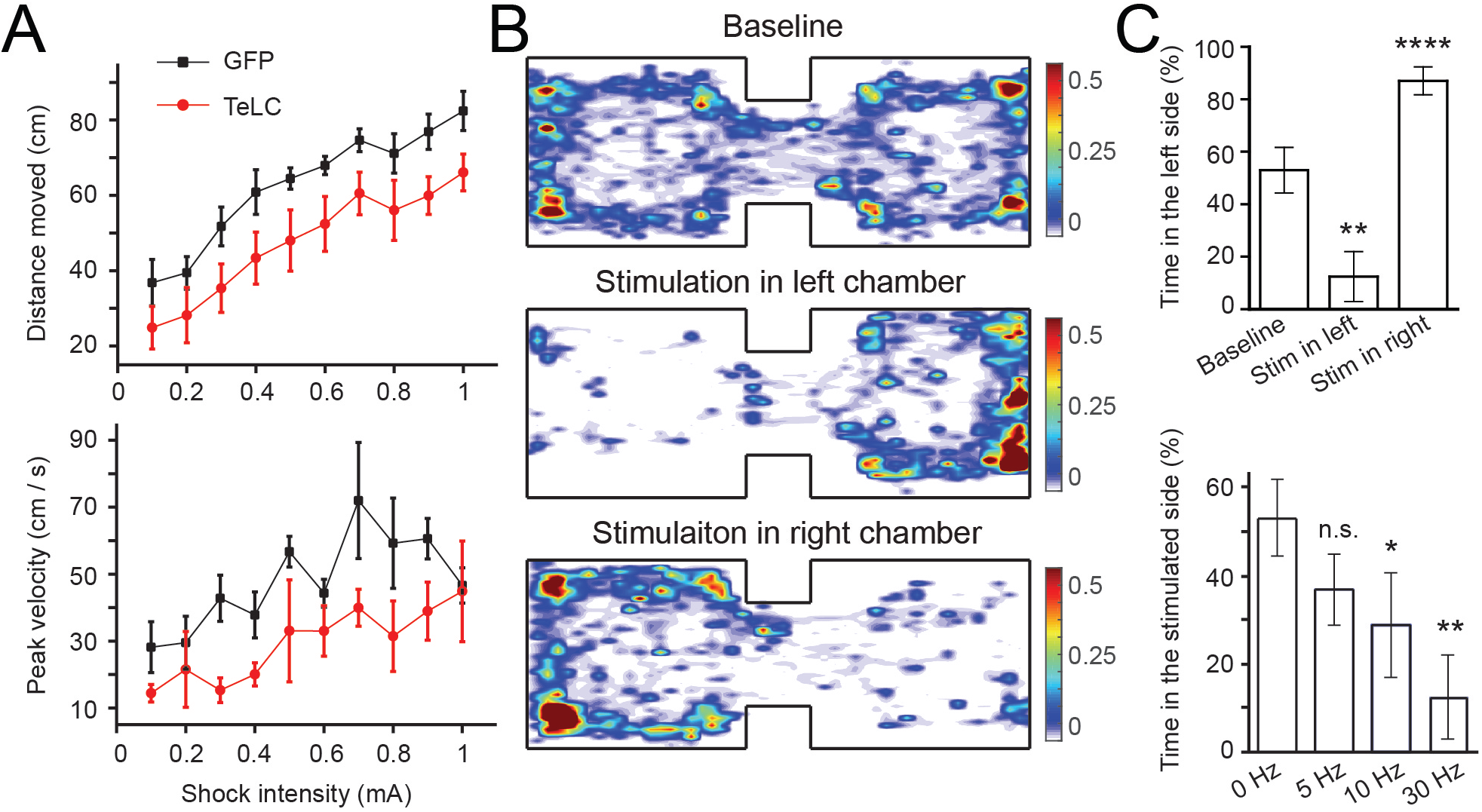
PKC-δ^+^ CeL neurons carry negative emotional valence. (A) Motor responses of mice within 2 seconds after foot shock (GFP, *Prkcd-Cre* mice in which PKC-δ^+^ CeL neurons expressed GFP, n = 4; TeLC, *Prkcd-Cre* mice in which PKC-δ^+^ CeL neurons expressed TeLC, n = 4; top panel, distance moved, F(1,6) = 13.6, P < 0.05; bottom panel, peak velocity, F(1,6) = 11.55, P < 0.05; two-way RM-ANOVA). (B) Heat-maps for the activity of a representative mouse at baseline (top), or during optogenetic activation of PKC-δ^+^ CeL neurons in either the left (middle) or right (bottom) chamber. (C) Top: the mice (n = 6) showed a significant place aversion to the chamber paired with laser activation of PKC-δ^+^ CeL neurons (F(1.9,9.5) = 104.5, P < 0.0001; **P < 0.01, ****P < 0.0001; one-way RM-ANOVA followed by Bonferroni’s test). Bottom: the place aversion effect is dependent on light stimulation frequency (F(2.1,10.3) = 21.48, P < 0.001; *P < 0.05, **P < 0.01, n.s., not significant (P > 0.05); one-way RM-ANOVA followed by Bonferroni’s test). Data are presented as mean ± s.e.m. in A and C.

**Fig. 8.**
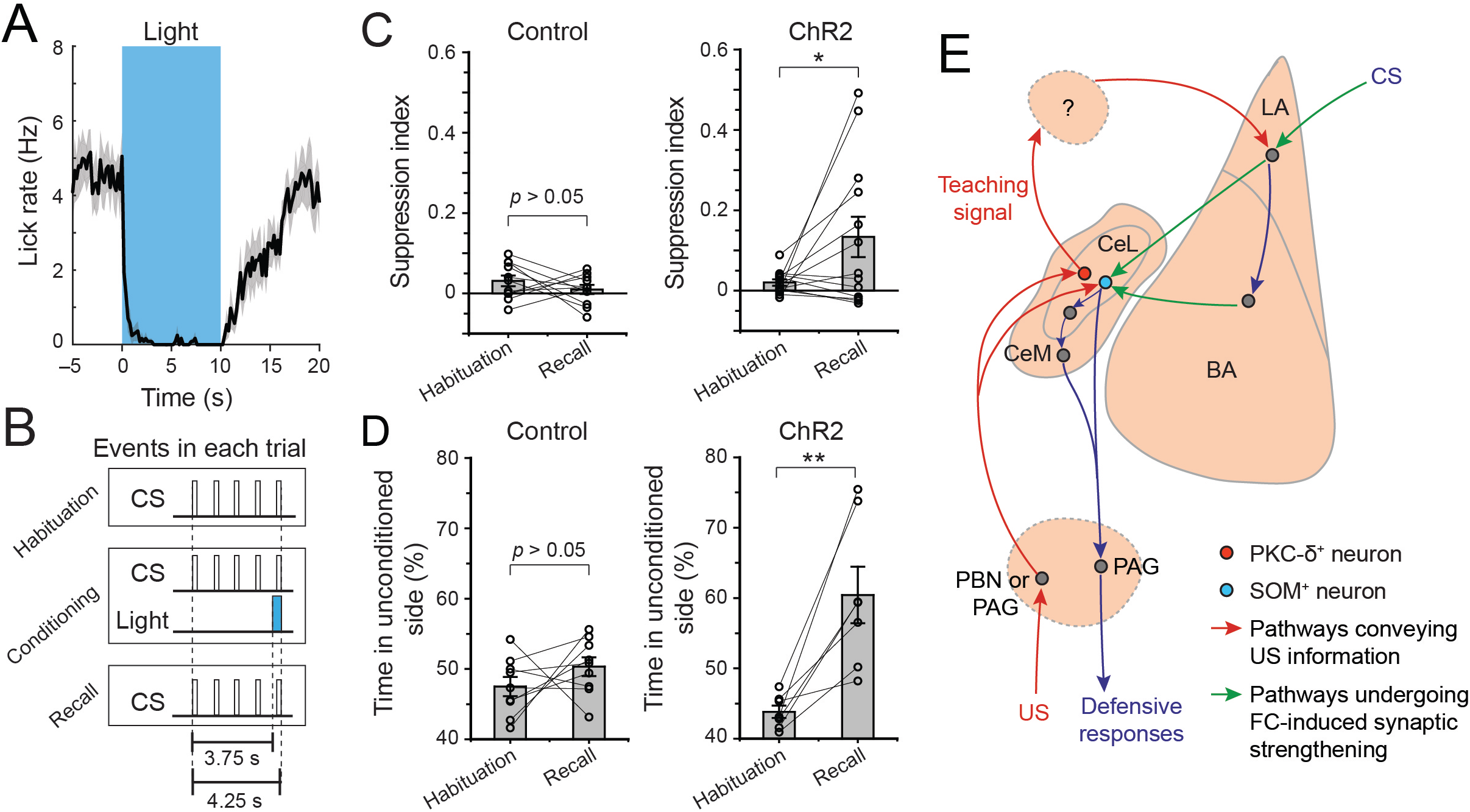
PKC-δ^+^ CeL neurons drive aversive learning. (A) Average lick rate of mice in which PKC-δ^+^ CeL neurons expressed ChR2. The blue bar denotes the time when light (473 nm, 30 Hz 5 ms pulses) was delivered to the CeL. Shaded area represents s.e.m. (B) A schematic of the experimental procedure. (C) Lick suppression, measured as suppression index, during baseline (habituation) and after conditioning (recall) by pairing sound with light stimulation of the CeL in mice in which PKC-δ^+^ CeL neurons expressed GFP (control) or ChR2 (control: n = 11, T(10) = 1, P > 0.05, paired t test; ChR2: n = 13, T(12) = 2.24, *P < 0.05, paired t test). (D) Place aversion during baseline (habituation) and after condition (recall) by pairing one side of a chamber with light stimulation of the CeL in mice in which PKC-δ^+^ CeL neurons expressed GFP (control) or ChR2 (control: n = 9, T(8) = 1.16, P > 0.05, paired t test; ChR2: n = 7, T(6) = 4.32, **P < 0.01, paired t test). (E) The proposed circuitry underlying classical fear conditioning. For clarity, only key known components of the circuitry are shown. Of note, both the PBN and the PAG may participate in providing the US signal during fear conditioning. As neurons in the CeL do not directly project to the LA, the question mark indicates structure(s) that may relay the teaching signal from the PKC-δ^+^ CeL neurons to the LA. Candidate structures include the SI or other areas innervated by PKC-δ^+^ CeL neurons. Data are presented as mean ± s.e.m. in A, C and D.

Furthermore, optogenetic activation of PKC-δ^+^ CeL neurons can be substituted for US to condition the mice for lick-suppression (Fig. 8B, C; Fig. S8B). PKC-δ^+^ CeL neuron activation appeared less efficient than tail shocks for the conditioning (see Fig. 3 & Fig. S3), likely because SOM^+^ CeL neuron activation is also required for learning and expression of the conditioned lick-suppression (Yu et al., 2016). We therefore tested whether PKC-δ^+^ CeL neuron activation could induce conditioned place aversion, a learned avoidance response that does not depend on SOM^+^ CeL neurons (see (Tan et al., 2012; Yu et al., 2016)). As expected, pairing optogenetic activation of PKC-δ^+^ CeL neurons with one side of a chamber readily caused mice to avoid that side when tested in the following day (Fig. 8D; Fig. S8B). Together, these results indicate that PKC-δ^+^ CeL neurons convey aversive information and are sufficient to drive aversive learning.

## DISCUSSION

Altogether, our results indicate that PKC-δ^+^ CeL neurons constitute a key node in a pathway that imparts information about US to the LA during FC, hence revealing a previously unknown amygdala functional organization, in which the CeL is upstream of the LA in processing aversive US during learning. In light of the current findings, together with recent discoveries that SOM^+^ CeL neurons are essential for both acquisition and expression of conditioned freezing behavior (Fig. 1D) (Li et al., 2013; Penzo et al., 2014; Penzo et al., 2015) and lick-suppression (Yu et al., 2016), we propose that the CeL participates in at least two distinct processes in FC (Fig. 8E): while PKC-δ^+^ CeL neurons convey the US information that serves as a “teaching signal” to instruct plasticity and learning in the LA, SOM^+^ CeL neurons undergo conditioning-induced synaptic plasticity that, together with the plasticity in the LA and elsewhere in the memory encoding network (Duvarci and Pare, 2014; Grundemann and Luthi, 2015; Herry and Johansen, 2014; LeDoux, 2000), enables the expression of learned defensive responses, in particular those of the passive type such as freezing behavior and suppression of action (Li et al., 2013; Penzo et al., 2014; Penzo et al., 2015; Yu et al., 2016).

The function of PKC-δ^+^ CeL neurons that we identify here may explain previous observations that the activities of some of the CeA neurons signal attention or salience (Calu et al., 2010; Roesch et al., 2012). Such function could be mediated by PKC-δ^+^ CeL neurons’ control over neuromodulatory systems. Indeed, these neurons project to and control the activity of different neuronal populations in the substantia innominata (SI) (data not shown; see also an accompanying paper in this issue of *Neuron* by Haohong Li and colleagues), a basal forebrain structure in which the cholinergic neurons have been implicated in attention and arousal (Hangya et al., 2015; Lin et al., 2015; Luchicchi et al., 2014). Of note, a recent study demonstrates that these cholinergic neurons send dense projections to the BLA, are critical for fear conditioning, and regulate BLA neuronal activity and synaptic plasticity (Jiang et al., 2016). It is therefore possible that SI cholinergic neurons may participate in carrying out the function of PKC-δ^+^ CeL neurons (Fig. 8E). Alternatively, or additionally, as PKC-δ^+^ CeL neurons have diverse projection targets (for reference, see an accompanying paper in this issue of *Neuron* by Haohong Li and colleagues), pathways other than that to the SI might mediate the function of these neurons.

Our findings also revise a prevalent model for the functional organization of amygdala circuits, which posits that PKC-δ^+^ CeL neurons are “fear-off” neurons – a CeL population that shows inhibitory CS responses following fear conditioning (Ciocchi et al., 2010) – that act to suppress fear responses through inhibition of amygdala output (Haubensak et al., 2010). In fact, we show that a substantial population of PKC-δ^+^ CeL neurons are essentially “fear-on” neurons and function in the opposite manner by conveying aversive US signals. These US signals likely come from the PBN or the periaqueductal gray (PAG) (Fig. 8E), in which neurons have been shown to provide either nociceptive or teaching signals, respectively, to the amygdala during fear conditioning (Han et al., 2015; Herry and Johansen, 2014; Johansen et al., 2010; Ozawa et al., 2017; Sato et al., 2015).

## AUTHOR CONTRIBUTIONS

K.Y. and B.L. designed the study. K.Y., S.A., X.Z., and H.S. conducted experiments (X.Z. performed the experiments in which LA neurons were imaged; S.A. and H.S. performed the experiments in which synaptic plasticity in LA neurons was examined; K.Y. conducted all the rest of the experiments). K.Y., X.Z., S.A., and H.S. analyzed data. C.R., L.F. and K.D. developed the intersectional viral strategy and provided critical reagents. P.Z. and L.P. developed and assisted with the imaging analysis methods (CNMF and CNMF-E). B.L. wrote the paper with inputs from all authors.

## ACKNOWLEDGMENTS

We thank Dr. Garret D. Stuber for helping with the *in vivo* imaging experiments. We thank Drs. Garret D. Stuber and Joshua Johansen for critically reading an earlier version of the manuscript, Ga-Ram Hwang for technical assistance, and members of the Li laboratory for helpful discussions. This work was supported by grants from the National Institutes of Health (NIH) (R01MH101214 to B.L.), NARSAD (to B.L, and S.A.), Louis Feil Trust (to B.L.), the Stanley Family Foundation (to B.L.), Simons Foundation (to B.L.), and Wodecroft Foundation (to B.L.).

## EXPERIMENTAL PROCEDURES

### Animals

Before surgery, mice were group-housed under a 12-h light-dark cycle (7 a.m. to 7 p.m. light) with food and water freely available. Animals with implants were housed singly. The *Som-cre* (Taniguchi et al., 2011), *Prkcd-Cre* (Haubensak et al., 2010), *Som-Flp* (Penzo et al., 2015), Ai32 and Ai35 (Madisen et al., 2012) mice have all been described elsewhere. The *Som-Cre* and *Som-Flp* mice were provided by Dr. Z. Josh Huang. The *Prkcd-Cre* mice were purchased from the Mutant Mouse Regional Resource Centers (MMRRC) as cryo-preserved spermatozoa (Donor: Dr. Nathaniel Heintz). Other mice were purchased from the Jackson Laboratory. All mice were bred onto C57BL/6J genetic background. The *Prkcd-Cre;Som-Flp* mice were bred by crossing the *Prkcd-Cre* mice with *Som-Flp* mice. Male and female mice of 40–60 d of age were used for all the experiments. All procedures involving animals were approved by the Institute Animal Care and Use Committees of Cold Spring Harbor Laboratory and carried out in accordance with US National Institutes of Health standards.

### Viral vectors

The AAV-DIO-TeLC-GFP, AAV-DIO-GFP, AAV-CAG-ChR2(H134R)-eYFP, AAV-DIO-ArchT-GFP, AAV-DIO-ChR2(H134R)-eYFP, and AAV-hSyn-GCaMP6f viruses (all serotype 2/9) were made by the Penn Vector Core (Philadelphia, PA). The AAV-hSyn-DIO-HA-KORD-IRES-mCitrine (2/8) virus was made by the University of North Carolina Vector Core (Chapel Hill, NC). The AAVdj-hSyn-C_on_/F_off_-GCaMP6m and AAVdj-hSyn-C_on_/F_off_-hChR2-mCherry viruses were made by the Stanford Vector Core (Stanford, CA). All viruses were stored in aliquots at –80 °C until use. We waited for at least 5 weeks after injection for optimal viral expression.

### Stereotaxic surgery

Standard surgical procedures were followed for stereotaxic injection (Li et al., 2013; Penzo et al., 2015). Briefly, mice were either anesthetized with ketamine (100 mg per kg of body weight) supplemented with dexmedetomidine hydrochloride (0.4 mg per kg), or anesthetized with isoflurane (using 2% at the beginning and 0.5–1% for the rest of the surgery procedure). Mice were positioned in a stereotaxic injection frame and laid on a heating pad maintained at 35°C. A digital mouse brain atlas was linked to the injection frame to guide the identification and targeting (Angle Two Stereotaxic System, myNeuroLab.com).

Viruses (∼0.3 μl) were delivered with a glass micropipette (tip diameter, ∼5 μm) through a skull window (1–2 mm^2^) by pressure applications (5–20 psi, 5–20 ms at 0.5 Hz) controlled by a Picrospritzer III (General Valve) and a pulse generator (Agilent). The injection was performed using the following stereotaxic coordinates for the CeL: –1.22 mm from Bregma, 2.9 mm lateral from the midline, and 4.6 mm vertical from skull surface; for the LA: –1.55 mm from Bregma, 3.2 mm lateral from the midline, and 4.2 mm vertical from skull surface; for the MGN: –3.16 mm from Bregma, 1.90 mm lateral from the midline, and 3.20 mm vertical from skull surface. For the *in vivo* photostimulation experiments, immediately after viral injection, mice were bilaterally implanted with optic fibers (core diameter, 105 μm; Thorlabs, Catalog number FG105UCA) that were placed above the CeL (coordinates of the fiber tip: –1.22 mm from Bregma, 2.9 mm lateral from the midline, and 4.3 mm vertical from skull surface). The optic fiber together with the ferrule (Thorlabs) was secured to the skull with C&B-Metabond Quick adhesive luting cement (Parkell Prod), followed by dental cement (Lang Dental Manufacturing).

For the *in vivo* imaging experiments, immediately after viral injection, a GRIN lens (diameter, 500 μm; ∼8.4mm length, part ID 130-000152; Inscopix) was implanted 200 μm above the center of injection.

Following the above procedures, a small piece of metal bar was mounted on the skull, which was used to hold the mouse in the head fixation frame during experiments.

### Immunohistochemistry

Immunohistochemistry experiments were performed following standard procedures. Briefly, mice were anesthetized with Euthasol (0.4 ml; Virbac, Fort Worth, Texas, USA) and transcardially perfused with 40 ml of PBS, followed by 40 ml of 4% paraformaldehyde in PBS. Brains were extracted and further fixed in 4% PFA overnight followed by cryoprotection in a 30% PBS-buffered sucrose solution for 36 h at 4 °C. Coronal sections (40 or 50 μm thickness) were cut using a freezing microtome (Leica SM 2010R, Leica). Sections were first washed in PBS (3 × 5 min), incubated in PBST (0.3% Triton X-100 in PBS) for 30 min at room temperature (RT) and then washed with PBS (3 × 5 min). Next, sections were blocked in 5% normal goat serum in PBST for 30 min at RT and then incubated with primary antibodies overnight at 4 °C. Sections were washed with PBS (5 × 15 min) and incubated with fluorescent secondary antibodies at RT for 1 h. After washing with PBS (5 × 15 min), sections were mounted onto slides with Fluoromount-G (eBioscience, San Diego, California, USA). Images were taken using a LSM 780 laser-scanning confocal microscope (Carl Zeiss, Oberkochen, Germany). The primary antibodies used were: rabbit anti-Somatostatin-14 (Peninsula Laboratories Inc., San Carlos, California, USA; catalogue number T-4103); mouse anti-PKC-δ (BD Biosciences, NJ, USA, cat. # 610397); rabbit anti-HA-Tag (C29F4, Cell Signaling, Danvers, MA, USA, cat. # 3724S). The fluorophore-conjugated secondary antibody used was Alexa Fluor^®^ 594 donkey anti-rabbit IgG (H+L) or Alexa Fluor^®^ 488 goat anti-rabbit IgG (H+L) (Life Technologies, Carlsbad, California, USA; catalogue number A21207 or A11008, respectively), depending on the desired fluorescence color.

### *In vitro* electrophysiology

To assess the synaptic plasticity in LA neurons induced by auditory fear conditioning, we specifically examined the synaptic transmission onto these neurons driven by the auditory thalamus – the medial geniculate nucleus (MGN) – that conveys the conditioned stimulus to the LA. To this end, we used mice in which the MGN was injected with the AAV-CAG-ChR2(H134R)-YFP, so that the MGN–LA pathway can be optogenetically stimulated in acute slices. Patch clamp recording was performed as previously described (Li et al., 2013; Penzo et al., 2015). Briefly, mice were anesthetized with isoflurane before they were decapitated; their brains were then dissected out and placed in ice chilled dissection buffer (110 mM choline chloride, 25 mM NaHCO3, 1.25 mM NaH2PO4, 2.5 mM KCl, 0.5 mM CaCl2, 7.0 mM MgCl2, 25.0 mM glucose, 11.6 mM ascorbic acid and 3.1 mM pyruvic acid, gassed with 95% O2 and 5% CO2). An HM650 Vibrating-blade Microtome (Thermo Fisher Scientific) was then used to cut 300 μm thick coronal sections that contained the amygdala. These slices were subsequently transferred to a storage chamber that contained oxygenated artificial cerebrospinal fluid (ACSF) (118 mM NaCl, 2.5 mM KCl, 26.2 mM NaHCO3, 1 mM NaH2PO4, 20 mM glucose, 2 mM MgCl2 and 2 mM CaCl2, at 34 °C, pH 7.4, gassed with 95% O2 and 5% CO2). Following 40 min of recovery, slices were transferred to RT (20-24 °C), where they were continuously bathed in the ACSF.

Visually guided whole-cell patch clamp recording from LA neurons was obtained with Multiclamp 700B amplifiers and pCLAMP 10 software (Molecular Devices, Sunnyvale, California, USA), and was guided using an Olympus BX51 microscope equipped with both transmitted and epifluorescence light sources (Olympus Corporation, Shinjuku, Tokyo, Japan). LA pyramidal neurons were identified for patching. Light-stimulation was used to evoke excitatory postsynaptic currents (EPSCs) driven by the ChR2-expressing axons originating from the MGN. The light source was a single-wavelength LED system (λ = 470 nm; CoolLED.com) connected to the epifluorescence port of the Olympus BX51 microscope. Light pulses of 0.2–0.5 ms, triggered by a TTL signal from the Clampex software (Molecular Devices), were used to evoke synaptic transmission. Synaptic responses were low-pass filtered at 1 KHz and recorded at holding potentials of –70 mV (for AMPA-receptor-mediated responses) and +40 mV (for NMDA-receptor-mediated responses). The AMPA/NMDA (A/N) ratio was calculated as the ratio of peak current at –70 mV to the current at 100 ms after light-stimulation onset at +40 mV (Clem and Huganir, 2010). Recordings were made in the ACSF with picrotoxin (100 μM) added. The internal solution contained 115 mM caesium methanesulphonate, 20 mM CsCl, 10 mM HEPES, 2.5 mM MgCl2, 4 mM Na2ATP, 0.4 mM Na3GTP, 10 mM sodium phosphocreatine and 0.6 mM EGTA (pH 7.2). The EPSCs were analysed using pCLAMP10 software (Molecular Devices).

### Behavioral tasks

#### Fear conditioning measuring conditioned freezing

We followed standard procedures for conventional auditory fear conditioning (Li et al., 2013; Penzo et al., 2014; Penzo et al., 2015). Briefly, mice were initially handled and habituated to a conditioning cage, which was a mouse test cage (18 cm × 18 cm × 30 cm) with an electrifiable floor connected to a H13-15 shock generator (Coulbourn Instruments). The test cage was located inside a sound attenuated cabinet (H10-24A; Coulbourn Instruments). Before each conditioning session the test cage was wiped clean with 70% ethanol. During conditioning the cabinet was illuminated and the behaviour was captured with a monochrome CCD-camera (Panasonic WV-BP334) at 3.7 Hz and stored on a personal computer. The FreezeFrame software (Coulbourn Instruments) was used to control the delivery of both tones and foot shocks. For habituation, five 4-kHz, 75-dB tones (conditioned stimulus), each of which was 30 s in duration, were delivered at variable intervals. During conditioning, mice received five presentations of the conditioned stimulus, each of which co-terminated with a 2-s, 0.7-mA foot shock (unconditioned stimulus). The recall of fear memory was tested 24 h following conditioning in a novel illuminated context, where mice were exposed to two presentations of unreinforced conditioned stimulus (120 s interstimulus interval). The novel context was a cage with a different shape (22 cm × 22 cm × 21 cm) and floor texture compared with the conditioning cage. Prior to each use the floor and walls of the cage were wiped clean with 0.5% acetic acid to make the scent distinct from that of the conditioning cage. Freezing responses to the conditioned stimuli were analyzed with FreezeFrame software (Coulbourn Instruments). The average of the freezing responses to the two conditioned stimuli during recall was used as an index of the conditioned fear.

#### Fear conditioning measuring conditioned lick-suppression

As previously described (Yu et al., 2016), water deprivation started 23 hours before training. Mice were trained to stay on a movable wheel under head fixation for 30 minutes in the first day, and 10 minutes daily afterwards. A metal spout was placed in front of animal mouth for water delivery. The spout also served as part of a custom “lickometer” circuit, which registered a lick event each time a mouse completed the circuit by licking the spout. The lick events were recorded by a computer through a custom software written in LabView (National Instruments). Each lick triggered a single opening of a water valve calibrated to deliver 0.3 μl water. It took mice 4–7 days to achieve stable licking, the criterion for which was 10-minute continuous lick without any gap longer than 10 s.

Mice with stable licking behavior were first subjected to sound habituation sessions (1 session / day for two days), during which auditory stimuli were presented through a computer speaker in each trial. Each stimulus was composed of 5 pips of pure tone (8 kHz, 70 dB). Pip duration was 250 ms, and inter-pip-interval was 750 ms. Each of the habituation sessions contained 15 trials with variable inter-trial-intervals (30-50 s). 24 h following habituation, mice were conditioned for 15 trials with variable inter-trial-intervals (30-50 s). In each of these trials, the auditory stimulus (CS) was presented and followed immediately by a tail shock (US; 100 μΑ for 500 ms), which was generated from an isolator (ISO-Flex, A.M.P. Instruments LTD, Israel) and delivered through a pair of wires secured to the tail with silicone tubing.

We used a lick suppression index to quantify animals’ performance in this task: Lick suppression index = (LPRE − LCS) / (LPRE + LCS), where LPRE is the number of licks in the 5 s period before CS onset, and LCS is the number of licks in the 5 s CS period (Yu et al., 2016).

#### Real time place aversion (RTPA)

As previously described (Stephenson-Jones et al., 2016), one side of a custom chamber (23 × 33 × 25 cm; made from plexiglass) was assigned as the stimulation zone, counterbalanced among mice. Mice were placed individually in the middle of the chamber at the onset of the experiment, the duration of which was 20 min. Laser stimulation (5-ms pulses delivered at 5, 10, or 30 Hz) was triggered when mice entered the stimulation zone, and lasted until mice exited the stimulation zone. Mice were videotaped with a CCD camera interfaced with the Ethovision software (Noldus Information Technologies), which was used to control the laser stimulation and extract the behavioral parameters (position, time, distance, and velocity).

#### Conditioned place aversion

The same chamber as that for the RTPA was used for the conditioned place aversion test. To make the two sides of the chamber distinct from each other, each side was decorated with a unique visual pattern (dotted vs striped) and scented with a unique odor (cherry vs. blueberry). The test consisted four sessions, each per day for four consecutive days. In session 1, the habituation session, mice were individually placed in the center of the chamber and allowed to freely explore both sides. In session 2 and 3, the conditioning sessions, the mice received 10 trials of laser stimulation, each consisting of a10-s train of 30-Hz 5-ms pulses, in one side of the chamber that was chosen as the stimulation side (counterbalanced among mice). The exit of the stimulation side was blocked during the conditioning. In session 4, the recall session, the mice were allowed to freely explore both sides of the chamber. The mice were videotaped in session 1 and 4 with a CCD camera interfaced with the Ethovision software (Noldus Information Technologies), which was used to control the laser stimulation and extract the behavioral parameters (position, time, distance, and velocity) (Stephenson-Jones et al., 2016).

### *In vivo* optogenetics

For bilateral optogenetic stimulation in the CeL in behaving mice, a rotary joint (Doric Lenses, Inc., Quebec, Canada, Catalog number FRJ_1x2i_FC-2M3_0.22) was used in the light delivery path, with one end of the rotary joint connected to a laser source (λ = 473 or 532 nm, OEM Laser Systems) and the other end, which has two terminals, to two optic fibers (for bilateral stimulation) through sleeves (Thorlabs). This configuration allows the mice carrying fiber-optic implants to freely move during optogenetic stimulation. The stimulation was typically composed of 5-ms 30-Hz light pulses delivered for various durations, unless otherwise specified. Laser intensity was 10 mW measured at the end of optic fibers.

### *In vivo* calcium imaging and analysis

We followed a recently described procedure for the *in vivo* imaging experiments (Resendez et al., 2016). All imaging experiments were conducted on awake behaving mice under head fixation in a dark, sound attenuated box. GCaMP6 fluorescence signals were imaged using a miniature, integrated fluorescence microscope system (Inscopix, Palo Alto, CA) with GRIN lenses implanted in the target areas (CeL and LA). We imaged PKC-δ^+^ CeL neuron activities while subjecting the mice to sound habituation, conditioning and recall sessions in the conditioned lick suppression task. Each session contained 15 trials, with random inter-trial-intervals (10-30 s). The same mice were subsequently used to image PKC-δ^+^ CeL responses to signaled and unsignaled shocks. A total of 16 shocks, 8 signaled and 8 unsignaled, were delivered in a randomly interleaved manner. In addition, the assignment of the first trial as having a signaled or unsignaled shock was counterbalanced among the mice.

We also imaged LA neuron responses to shocks before and after subcutaneous injection (s.c.) of SALB (10 mg / kg of body weight) (to inhibit PKC-δ^+^ CeL neurons). As a control experiment, we imaged LA neuron responses to shocks before and after systemic application (s.c.) of DMSO (the vehicle of SALB). Each session contained 15 trials.

For the experiments in which PKC-δ^+^ CeL neurons were imaged, we installed a baseplate on top of the GRIN lens for each mouse, as described previously (Resendez et al., 2016). Before imaging, the miniature microscope was attached to the baseplate. The microscope was adjusted such that the best dynamic fluorescence signals were at the focal plane, which was subsequently kept constant across imaging sessions. For the experiments in which LA neurons were imaged, the microscope was mounted on top of, and aligned with the GRIN lens with a custom adjustable micromanipulator that allows movement in all 3 axes. The focus of the microscope was adjusted through the micromanipulator to get the best focal plane as described above.

During imaging, the Data Acquisition Box of the nVista Imaging System (Inscopix, Palo Alto, CA) was triggered by an NI data acquisition device (USB6008, National Instruments, CA). Compressed gray scale images were then recorded with nVistaHDV2 (Inscopix) at 10 frames per second. The analog gain (1 to 5) and LED output power (8% to 30% of the maximum) of the microscope were set to be constant for the same subject across imaging sessions. During imaging, the time stamps of different events, including the trigger signals sent to the microscope, the auditory stimuli, the electrical shocks, and the licks, were recorded with a custom program written in LabView (National Instruments, CA).

For imaging data processing and analysis, we began by importing the compressed video files into Mosaic (version 1.0.0b; Inscopix, Palo Alto, CA), in which we trimmed the first frame of the video for each trial to minimize the influence of the flash associated with LED light onset. We subsequently used Mosaic to combine all the trimmed video files into a single .tiff stack, apply a 4-pixel bin to the .tiff stack, and correct the motion artifact. A new .tiff stack was then saved for further processing.

Next, to address the problem of high levels of background fluorescence signals intrinsic to one-photon imaging, we applied our newly developed imaging analysis method, the extended Constrained Nonnegative Matrix Factorization (CNMF-E) (Zhou et al., 2016), in which we model the background with two realistic components: (1) one models the constant baseline and slow trends of each pixel, and (2) the other models the fast fluctuations from out-of-focus signals and is therefore constrained to have low spatial-frequency structure. This decomposition avoids cellular signals being absorbed into the background term. After subtracting the background approximated with this model, we used Constrained Nonnegative Matrix Factorization (CNMF) (Pnevmatikakis et al., 2016) to demix neural signals and get their denoised and deconvolved temporal activity, termed as ΔF. The CNMF-E method was carried out using a custom Matlab algorithm (for a detailed description of this method, see (Zhou et al., 2016)).

Once the temporal activity of the neurons was extracted, we characterized the CS (sound) or US (shock) responses of each neuron using auROC (area under the receiver-operating characteristic curve) analysis, in which we compared the average ΔF values during the baseline period (2 s immediately prior to the delivery of CS or US) with that during CS or US presentations (2 s immediately after the onset of CS or US) by moving a criterion from zero to the maximum ΔF value. We then plotted the probability that the ΔF values during CS or US presentations was greater than the criteria against the probability that the baseline ΔF values was greater than the criteria. The area under this curve quantifies the degree of overlap between the two ΔF distributions (i.e., the discriminability of the two). A permutation test (iteration 5000 times) was used to determine whether the average ΔF values during CS or US presentations were significantly (P < 0.05) higher than during baseline, and thus classify a neuron as being CS-responsive or US-responsive, respectively. The peak CS or US response amplitude in each trail was determined by searching the maximum value within a 5-s time window immediately after stimulus onset.

### Statistics and data presentation

All statistics are indicated where used. Statistic analyses were performed with Origin8 Software (OriginLab Corporation, Northampton, MA), GraphPad Prism Software (GraphPad Software, Inc., La Jolla, CA), or Matlab (MathWorks, Natick, MA). All behavioral experiments were controlled by computer systems, and data were collected and analyzed in an automated and unbiased way. Virus-injected animals in which the injection site was incorrect were excluded. For in vivo imaging, a mouse was excluded if the tract and tip of the GRIN lens was outside of the targeted area. No other mice or data points were excluded.

